# The medial temporal lobe is critical for spatial relational perception

**DOI:** 10.1101/765222

**Authors:** Nicholas A. Ruiz, Michael R. Meager, Sachin Agarwal, Mariam Aly

## Abstract

The medial temporal lobe (MTL) is traditionally considered to be a system that is specialized for long-term memory. Recent work has challenged this notion by demonstrating that this region can contribute to many domains of cognition beyond long-term memory, including perception and attention. One potential reason why the MTL (and hippocampus specifically) contributes broadly to cognition is that it contains relational representations — representations of multidimensional features of experience and their unique relationship to one another — that are useful in many different cognitive domains. Here, we explore the hypothesis that the hippocampus/MTL plays a critical role in attention and perception via relational representations. We compared human participants with MTL damage to healthy age- and education-matched individuals on attention tasks that varied in relational processing demands. On each trial, participants viewed two images (rooms with paintings). On ‘*similar room*’ trials, they judged whether the rooms had the same spatial layout from a different perspective. On ‘*similar art*’ trials, they judged whether the paintings could have been painted by the same artist. On ‘*identical*’ trials, participants simply had to detect identical paintings or rooms. Patients were significantly and selectively impaired on the similar room task. This work provides further evidence that the hippocampus/MTL plays a ubiquitous role in cognition by virtue of its relational and spatial representations, and highlights its important contributions to rapid perceptual processes that benefit from attention.

## Introduction

The human medial temporal lobe (MTL) has traditionally been considered a “memory system” of the brain — a system specialized for the formation and retention of long-term declarative memories (Cohen & Eichenbaum, 1993; Eichenbaum & Cohen, 2001; Milner et al., 1998; Squire & Wixted, 2011; Suzuki, 2009). Such a view has been challenged in recent years, with the emergence of a large body of work highlighting the many ways that the MTL, including the hippocampus, contributes to cognition beyond long-term declarative memory (Aly & Turk-Browne, 2018; Nadel & Peterson, 2013; Olsen et al., 2012; Shohamy & Turk-Browne, 2013). This includes studies showing a role for the hippocampus/MTL in perception (Lee et al., 2012), working memory (Yonelinas, 2013), implicit memory (Hannula & Greene, 2012), decision making (Shohamy & Daw, 2015), imagination (Schacter et al., 2017), creativity (Rubin et al., 2015), language (Duff & Brown-Schmidt, 2012), and social cognition (Schafer & Schiller, 2018).

In addition to this expanding literature, we have recently discovered that the hippocampus is involved in online attention behavior (Aly & Turk-Browne 2016a; Aly & Turk-Browne 2016b; Cordova, Turk-Browne, & Aly, 2019); It exhibits distinct activity patterns for different attentional states, and the stability of these activity patterns predicts attentional performance. However, this work implicating the human hippocampus in attention comes from fMRI studies, which do not tell us if the hippocampus contributes a *necessary* function for attention behavior. Here, we sought to determine whether the hippocampus (and MTL more broadly) plays a critical role in attention and to elucidate the nature of its contribution.

To that end, we tested patients with hippocampal/MTL damage on a modified version of the “art gallery” task we used previously, in fMRI studies, to demonstrate hippocampal involvement in attention (Aly & Turk-Browne 2016a; Aly & Turk-Browne, 2016b). This task allowed us to test two complementary hypotheses about how MTL function might support attention. Given the extensive literature demonstrating that a key aspect of hippocampal function is its *relational representations* (representations that link multiple features of an experience and their unique relationship to one another; Eichenbaum & Cohen, 2014; Konkel & Cohen, 2009; Olsen et al., 2012), one prediction is that patients with hippocampal damage will be impaired on tasks that require attention to the *relations* between features (e.g., Cordova et al., 2019). An alternative hypothesis is that hippocampal damage may only impair attention behaviors that require *spatial* representations. Such a prediction would be consistent with our previous fMRI work, in which the hippocampus was more strongly modulated by, and predicted behavior more strongly for, attention tasks that put demands on processing spatial layouts vs. artistic features (Aly & Turk-Browne 2016a; Aly & Turk-Browne, 2016b). Likewise, in our previous studies, MTL cortical regions were also more strongly modulated by attention to spatial relations (Aly & Turk-Browne 2016a; Aly & Turk-Browne, 2016b). This prediction is consistent with models of hippocampal function that emphasize its importance for spatial cognition (Maguire & Mullally, 2013; O’Keefe & Nadel, 1978), as well as findings that hippocampal damage impairs perception of complex scenes but not complex objects (Lee et al., 2005a, 2005b, 2012; in contrast, perirhinal cortex damage can impair perception of objects).

To test these hypotheses, we designed a task that required different kinds of attention across different trials that utilized the same type of stimulus (3D-rendered rooms with paintings). Trials varied in whether they placed a heavy demand on relational processing or a lighter demand. They also varied in whether they required attention to spatial features or object features.

This task has several important components. First, this approach allows us to determine how the hippocampus/MTL contribute to goal-directed attention when bottom-up stimulation is held constant because the same type of stimulus is used across trials in which participants’ behavioral goals are different. Second, stimuli were briefly presented and trial-unique, so that long-term memory was neither required nor beneficial for task performance (e.g., Aly et al., 2013). Finally, trials that placed heavy demands on relational processing could not be performed accurately by attending only to low-level visual features: instead, attending to abstract relations was essential (e.g., Hartley et al., 2007).

Together, these task features enabled us to rigorously test alternative theories of whether and how the hippocampus/MTL might critically contribute to attention and perception.

## Materials and Methods

### Participants

#### Demographics and Recruitment

Patients with medial temporal lobe lesions (n = 7; 1 woman, 6 men; *M_age_*= 41.0 years, *M_education_*= 17.0 years) were recruited via the New York University Patient Registry for the Study of Perception, Emotion, and Cognition (NYU PROSPEC) and the Department of Neurology at Columbia University Irving Medical Center. These patients had normal or corrected-to-normal vision and normal hearing. Neuropsychological test scores are shown in **Table 1** and **Table 2**.

**Table 1.** Neuropsychological examination scores for the hypoxia patient (101). The maximum raw score on the Montreal Cognitive Assessment (MoCA) is 30; 26 is considered the standard cutoff for identifying impairment. The MoCA section associated with the largest deduction of points was the delayed-recall trial of the memory test (0/5). Scores for the Repeatable Battery for the Assessment of Neuropsychological Status (RBANS) are adjusted for age. RBANS Standard Scores (SS) are based upon a mean of 100 and a standard deviation of 15.

| Neuropsychological Exam | Patient 101 |
| --- | --- |
| *Montreal Cognitive Assessment* | 21 |
| *Repeatable Battery for the Assessment of Neuropsychological Status* | 67 |
| Immediate Memory | 58 |
| Visuospatial | 115 |
| Language | 75 |
| Attention | 73 |
| Delayed Memory | 49 |

**Table 2.**
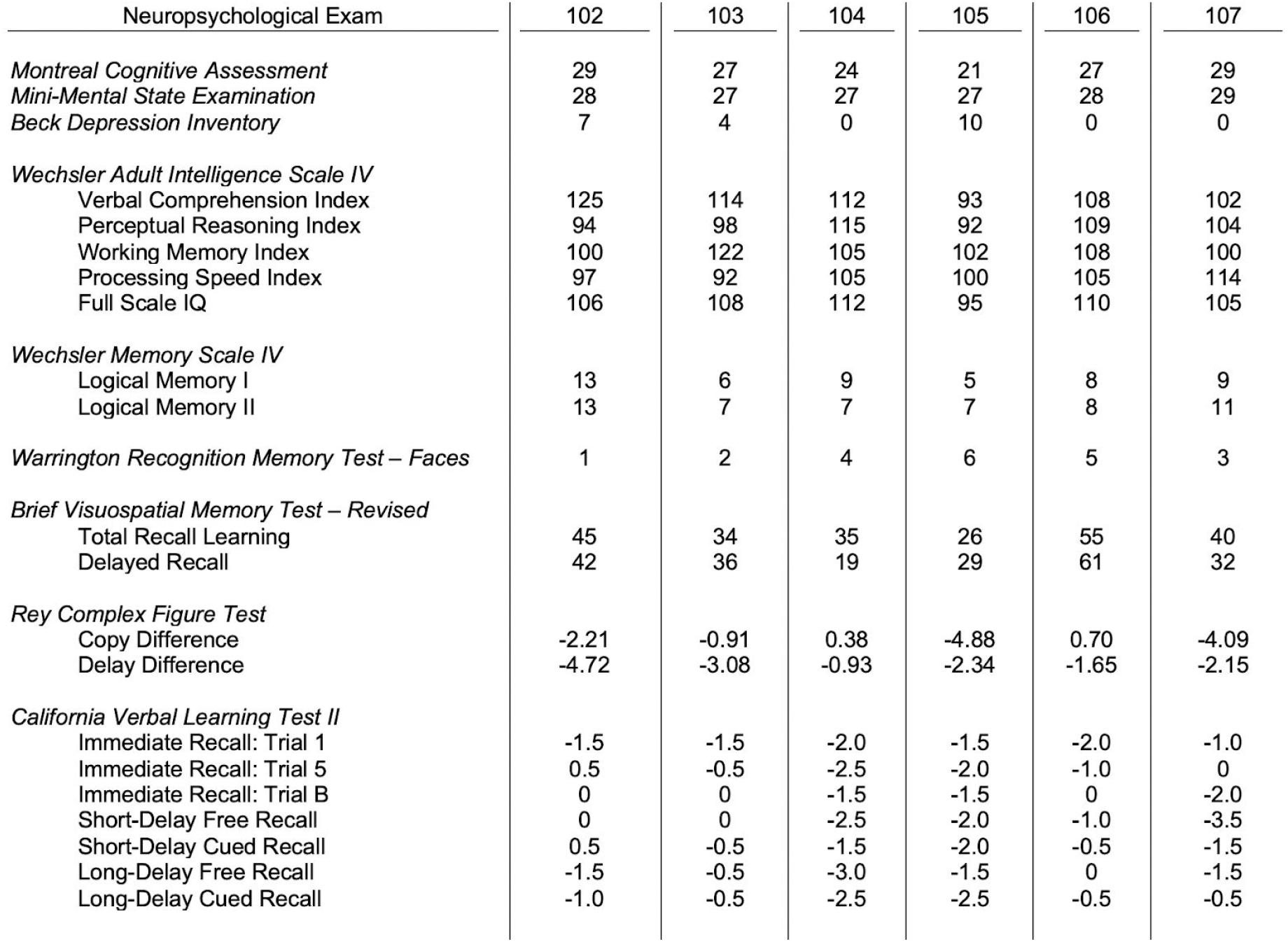
Neuropsychological examination scores for temporal lobectomy patients (102 - 107). The maximum raw score on the Montreal Cognitive Assessment (MoCA) is 30; a score of 26 is considered the standard cutoff for identifying impairment. The MoCA section associated with the largest deduction of points for all patients was the delayed-recall trial of the memory test. The maximum score on the Mini-Mental State Examination is 30, with the standard cutoff for identifying impairment being 24. Raw scores below 10 on the Beck Depression Inventory indicate a minimal level of depressive symptomatology characteristic of the general population. The index scores for the Wechsler Adult Intelligence Scale-Fourth Edition (WAIS-IV) are represented in Standard Scores (SS), which are based upon a mean of 100 and a standard deviation of 15. The Logical Memory subtest of the Wechsler Memory Scale-Fourth Edition (WMS-IV) and the Faces subtest of the Warrington Recognition Memory Test (WRMT) scores are adjusted for age; yielding scaled scores (ss) based upon a mean of 10 and a standard deviation of 3. The Brief Visuospatial Memory Test-Revised (BVMT-R) yields age corrected T-scores, which have a mean of 50 and a standard deviation of 10. The Rey Complex Figure Test (RCFT) scores are z-scores that are corrected for participant age, based upon a mean of 0.0 and a standard deviation of 1.0. The California Verbal Learning Test - 2nd Edition (CVLT-II) scores are normative z-scores corrected for age, based upon a mean of 0.0 and a standard deviation of 1.0.

| Neuropsychological Exam | 102 | 103 | 104 | 105 | 106 | 107 |
| --- | --- | --- | --- | --- | --- | --- |
| *Montreal Cognitive Assessment* | 29 | 27 | 24 | 21 | 27 | 29 |
| *Mini-Mental State Examination* | 28 | 27 | 27 | 27 | 28 | 29 |
| *Beck Depression Inventory* | 7 | 4 | 0 | 10 | 0 | 0 |
| *Wechsler Adult Intelligence Scale IV* |  |  |  |  |  |  |
| Verbal Comprehension Index | 125 | 114 | 112 | 93 | 108 | 102 |
| Perceptual Reasoning Index | 94 | 98 | 115 | 92 | 109 | 104 |
| Working Memory Index | 100 | 122 | 105 | 102 | 108 | 100 |
| Processing Speed Index | 97 | 92 | 105 | 100 | 105 | 114 |
| Full Scale IQ | 106 | 108 | 112 | 95 | 110 | 105 |
| *Wechsler Memory Scale IV* |  |  |  |  |  |  |
| Logical Memory I | 13 | 6 | 9 | 5 | 8 | 9 |
| Logical Memory II | 13 | 7 | 7 | 7 | 8 | 11 |
| *Warrington Recognition Memory Test – Faces* | 1 | 2 | 4 | 6 | 5 | 3 |
| *Brief Visuospatial Memory Test – Revised* |  |  |  |  |  |  |
| Total Recall Learning | 45 | 34 | 35 | 26 | 55 | 40 |
| Delayed Recall | 42 | 36 | 19 | 29 | 61 | 32 |
| *Rey Complex Figure Test* |  |  |  |  |  |  |
| Copy Difference | -2.21 | -0.91 | 0.38 | -4.88 | 0.70 | -4.09 |
| Delay Difference | -4.72 | -3.08 | -0.93 | -2.34 | -1.65 | -2.15 |
| *California Verbal Learning Test II* |  |  |  |  |  |  |
| Immediate Recall: Trial 1 | -1.5 | -1.5 | -2.0 | -1.5 | -2.0 | -1.0 |
| Immediate Recall: Trial 5 | 0.5 | -0.5 | -2.5 | -2.0 | -1.0 | 0 |
| Immediate Recall: Trial B | 0 | 0 | -1.5 | -1.5 | 0 | -2.0 |
| Short-Delay Free Recall | 0 | 0 | -2.5 | -2.0 | -1.0 | -3.5 |
| Short-Delay Cued Recall | 0.5 | -0.5 | -1.5 | -2.0 | -0.5 | -1.5 |
| Long-Delay Free Recall | -1.5 | -0.5 | -3.0 | -1.5 | 0 | -1.5 |
| Long-Delay Cued Recall | -1.0 | -0.5 | -2.5 | -2.5 | -0.5 | -0.5 |

Healthy adults (n = 14; 9 women, 5 men; *M_age_*= 42.0 years*, M_education_*= 15.8 years) were recruited via flyers posted around the Columbia University community. These participants reported no neurological or psychiatric illness, had normal or corrected-to-normal vision, and normal hearing. 12 of these individuals completed brief neuropsychological assessments. They scored in the normal range on the Montreal Cognitive Assessment (*M* = 28.36, *SD* = 1.22; max score = 30, 26+ is normal), the Mini-Mental State Examination (*M* = 29.00, *SD* = 1.41; max score = 30, 24+ is normal), and the Beck Depression Inventory (*M* = 4.50, *SD* = 3.71; scores below 10 are considered normal; one individual scored 13, indicating a mild mood disturbance).

Patients and healthy adults did not differ in age (*t*_19_= −0.12, *p* = 0.91, 95% CIs = −18.72 - 16.72) or education (*t*_19_= 0.93, *p* = 0.37, 95% CIs = −1.53 - 3.96).

#### Patient Descriptions

Patient 101 was recruited from the Columbia University Irving Medical Center. He suffered a hypoxic brain injury as a result of a period of asphyxiation and suspected cardiac arrest. An initial MRI revealed no signs of volume loss in the brain, but a follow-up FLAIR scan one year after the hypoxic event revealed hippocampal abnormalities. This was confirmed by a volumetric lesion analysis (see *Lesion Analyses* and **Table 3**).

**Table 3.** Volumetric analysis of the hippocampus and amygdala for the hypoxia patient (patient 101) and healthy age-matched participants (n=3). Values are region-of-interest volumes divided by total intracranial volume, to correct for differences in overall head and brain size. Z-scores < −1.96 (in bold) indicate statistically significant volume reductions (*p* <.05). SD = standard deviation.

|  | left hippocampus | right hippocampus | left amygdala | right amygdala |
| --- | --- | --- | --- | --- |
| patient 101 | 0.00207 | 0.00187 | 0.00093 | 0.00073 |
| control mean | 0.00259 | 0.00270 | 0.00100 | 0.00097 |
| control SD | 0.00013 | 0.00017 | 0.00009 | 0.00014 |
| patient z-score | **-4.16** | **-5.00** | -0.75 | -1.78 |

Relatively selective volume loss in the hippocampus is sometimes observed for mild hypoxia (Gadian et al., 2000; Hopkins et al., 1995; Rempel-Clower et al., 1996; Smith et al., 1984). However, volume reductions have also been observed in other subcortical structures (Guderian et al., 2015; Huang & Castillo, 2008). As a result, we cannot rule out brain damage outside of the hippocampus for patient 101. Indeed, although not rigorously analyzed, we believe his scans show some evidence of striatal and thalamic abnormalities. We include this patient because of the observed hippocampal abnormalities and volume reduction, but do not claim that the lesion is selective to the hippocampus.

Patients 102 - 107 were recruited via NYU PROSPEC. They underwent unilateral temporal lobectomies for the treatment of intractable epilepsy. The surgeries were to the right hemisphere for patients 102 and 103 and to the left hemisphere for the remaining patients. Because of the small sample size, we did not statistically compare the behavior of left and right hemisphere lesion patients, although laterality may affect performance (Willment & Golby, 2013).

The resected tissue in these patients was primarily the anterior hippocampus and surrounding medial temporal lobe cortex, as well as some sections of the lateral temporal cortex. Patient 106 additionally had a partial frontal lobe resection. Lesion masks are shown for 5 of the 6 temporal lobectomy patients in **Figure 1A**. Regions of lesion overlap include the anterior hippocampus and medial temporal lobe cortex.

**Figure 1.**
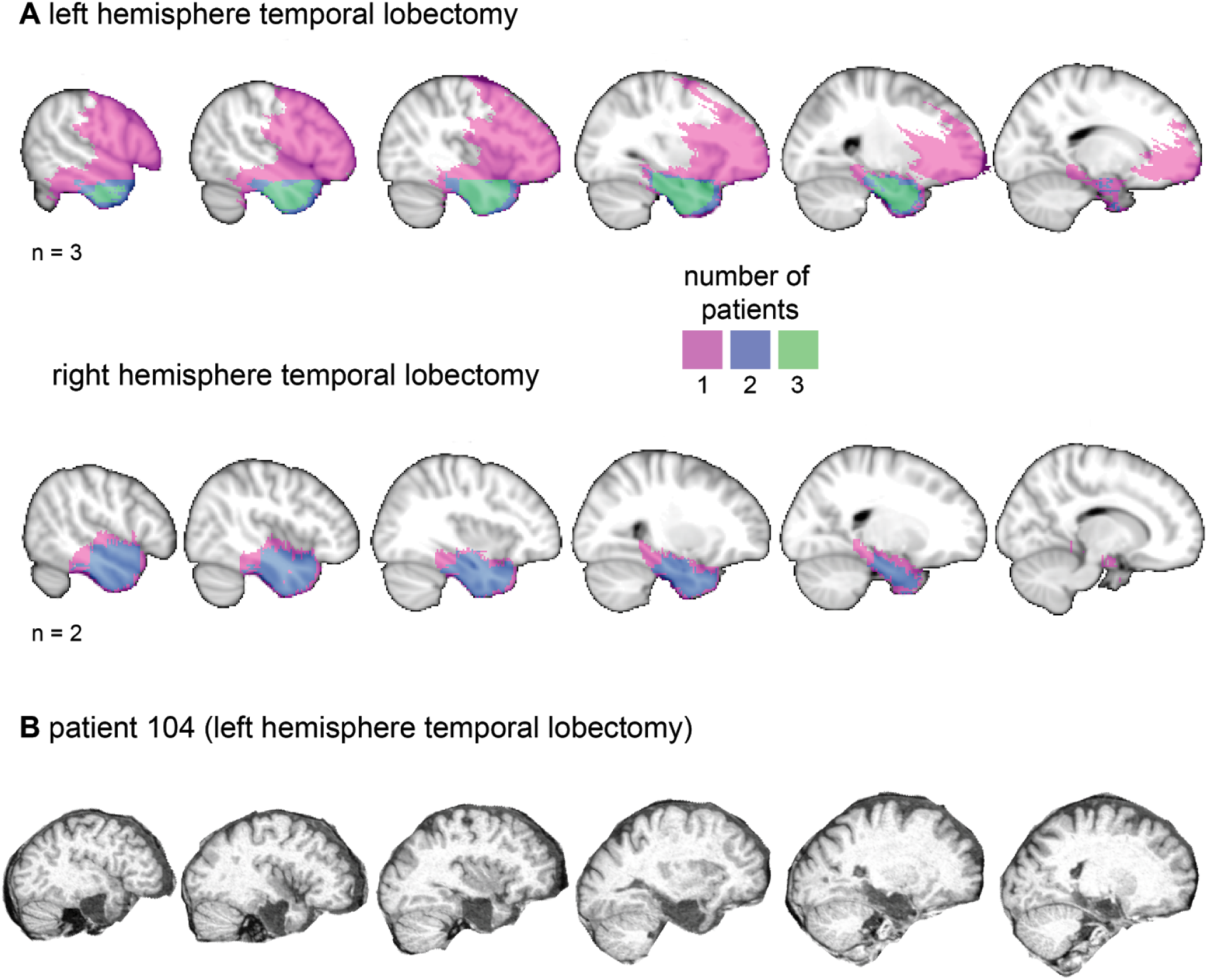
**(A)**Lesion masks for 5 (out of 6) patients that underwent unilateral temporal lobectomies. Colors indicate the number of patients with a lesion in a given location. All patients had damage to the hippocampus and surrounding medial temporal lobe cortex, with maximum overlap in the anterior hippocampus and anterior medial temporal lobe cortex. **(B)** Representative slices from the MRI of the left temporal lobectomy patient who did not have a lesion mask. This patient’s damage included the hippocampus and medial temporal lobe cortex.

Lesion masks were not available for one left temporal lobectomy patient (104), so representative slices from his MRI scan are shown in **Figure 1B**. This patient also had damage to the anterior hippocampus and medial temporal lobe cortex.

Thus, all of the patients — whether their etiology was hypoxia or temporal lobe epilepsy — had hippocampal/MTL atrophy. The damage outside of the MTL varied across patients; thus, we can infer that any resulting behavioral changes can be attributed to direct or indirect effects of MTL damage.

The study was approved by the Columbia University Institutional Review Board. All participants received monetary compensation ($15/hour for the experiment and for travel time). They gave written informed consent, filled out a demographics form, and completed neuropsychological examinations. These included the Montreal Cognitive Assessment (MoCA; Nasreddine et al., 2005), the Mini-Mental State Examination (MMSE; Folstein et al., 1975), and Beck Depression Inventory (BDI; Beck et al., 1961). The temporal lobectomy patients additionally completed the Wechsler Adult Intelligence Scale-Fourth Edition (WAIS-IV; Wechsler, 2008), the Logical Memory subtest of the Wechsler Memory Scale-Fourth Edition (WMS IV; Wechsler, 2009), the Warrington Recognition Memory Test – Faces Subtest (RMF; Warrington, 1996), the Brief Visuospatial Memory Test-Revised (BVMT-R; Benedict et al., 1996), the Rey Complex Figure Test (RCFT; Osterrieth, 1944; Rey, 1941), and the California Verbal Learning Test-Second Edition (CVLT-II; Delis et al., 2000) as part of a separate, extended neuropsychological evaluation conducted by neuropsychologists through NYU PROSPEC. The hypoxia patient completed the Repeatable Battery for the Assessment of Neuropsychological Status (RBANS; Randolph, Tierney, Mohr, & Chase, 1998) as part of a separate neuropsychological evaluation at the Columbia University Irving Medical Center. The hypoxia patient and temporal lobectomy patients completed different neuropsychological tests because they were assessed at different hospitals that had different norms for which tests are conducted. Unfortunately, the hypoxia patient moved across the country shortly after testing and no longer wishes to participate in research studies; thus, we are unable to perform further neuropsychological testing with him.

### Lesion Analyses

#### Temporal Lobectomy Patients

Patient MRIs were first transformed to MNI standard space. Then two neuropsychologists manually traced the lesion masks in FSLview, using a semi-transparent overlap of three images (MNI standard brain, patient’s MRI, lesion mask). FLAIR images were consulted in an adjacent window to provide additional cues with respect to lesion extent. Lesion masks were traced in 3 planes (coronal, sagittal, horizontal) and then reviewed again for corrections. Lesion masks were available for 5 (out of 6) temporal lobectomy patients **(Figure 1)**.

#### Hypoxic Patient

Volumetric analyses were conducted for patient 101 to characterize the nature and extent of his hippocampal damage **(Table 3)**. Hippocampal and amygdala volumes were compared for this patient and three age- and education-matched adults (patient 101; 19 years old, 12 years of education; healthy adults: *M_age_* = 18, *M_education_* = 13.7). First, Freesurfer was used to obtain intensity-normalized MRIs (via autorecon1). These images were then converted to NIFTI for use in an FSL (https://fsl.fmrib.ox.ac.uk) pipeline. After brain extraction, FIRST (Patenaude et al., 2011) was applied to automatically segment subcortical structures, including the hippocampus and amygdala. Hippocampal and amygdala segmentations were then manually edited by expert raters to correct imperfections from the automated approach. FAST (Zhang et al., 2001) was then used to obtain gray matter, white matter, and cerebrospinal fluid masks. These were summed in order to measure total intracranial volume. The volumes of the left and right hippocampus and amygdala were then divided by the total intracranial volume to correct for differences in overall head and brain size. These analyses revealed that the hippocampus, but not the amygdala, exhibited significant volume reductions in the hypoxia patient. Nevertheless, we cannot rule out brain damage elsewhere, particularly in the striatum and thalamus (Guderian et al., 2015; Huang & Castillo, 2008).

### Stimuli

Participants viewed images of rooms with paintings **(Figure 2)**. The rooms each contained multiple pieces of furniture, diverse wall angles, and a single painting. The paintings were primarily of outdoor scenes and spaces; some paintings also contained people. A subset of these stimuli has previously been used (Aly & Turk-Browne 2016a; Aly & Turk-Browne, 2016b).

**Figure 2.**
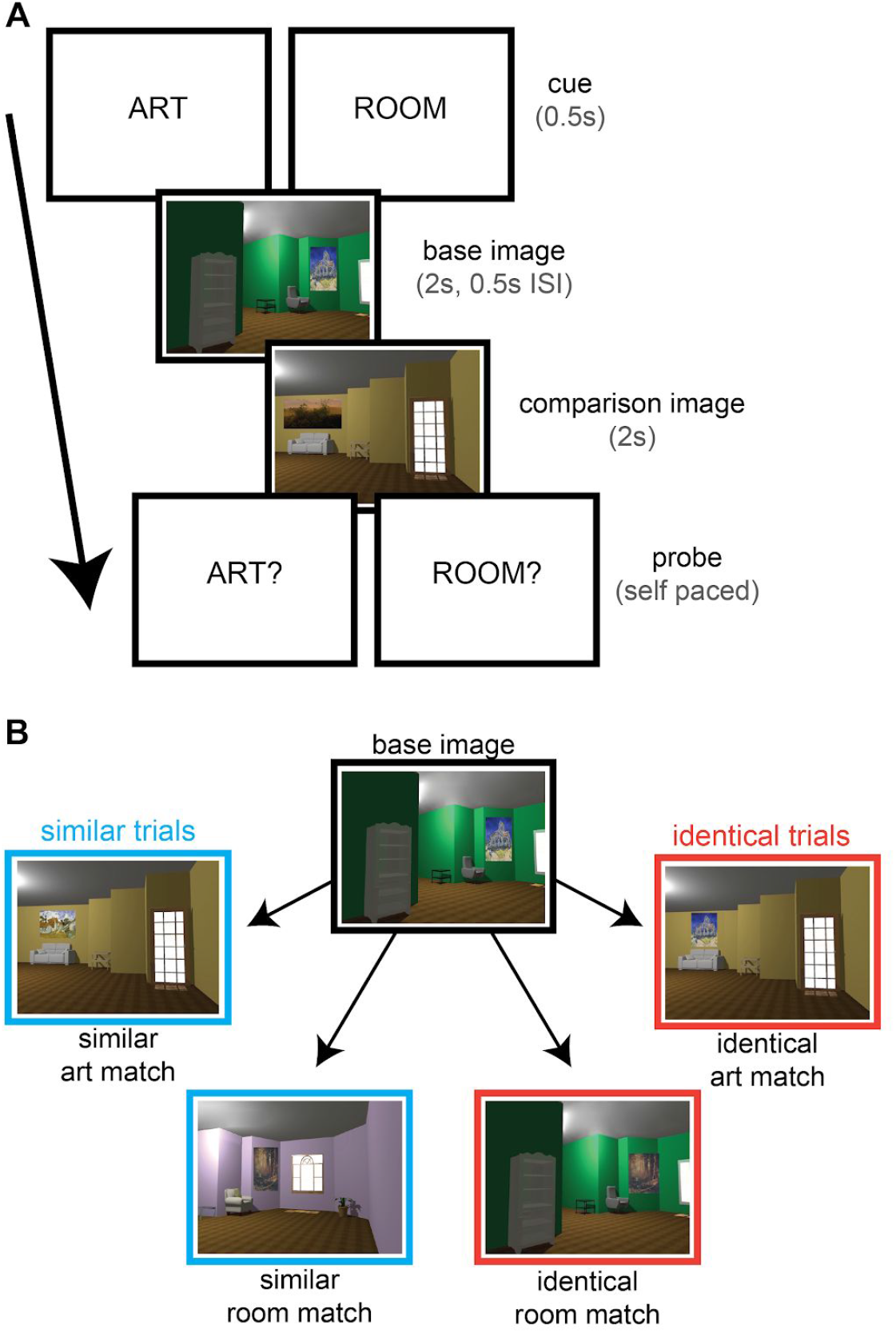
**(A)** Trial structure. Participants viewed two images on each trial. Prior to trial onset, they were instructed to attend to either the style of the paintings (ART) or the layout of the rooms (ROOM). At the end of the trial, participants were asked if the two paintings matched (ART?) or if the two rooms matched (ROOM?). An art match could be either two identical paintings (identical art match) or two different paintings by the same artist (similar art match). A room match could be either two identical rooms (identical room match) or two rooms with the same spatial layout from a different perspective (similar room match). On valid trials, the cue at the beginning of the trial was the same as the probe at the end; on invalid trials, the cue and probe were different. **(B)** Examples of a similar art match, similar room match, identical art match, and identical room match. A non-matching image (neither an art nor a room match) could also be displayed as the comparison image, as in **(A)**. Identical and similar trials were intermixed so that participants could not adopt different strategies during the viewing of the first image. ISI = inter-stimulus interval.

Rooms were created in Sweet Home 3D (http://www.sweethome3d.com/). 80 rooms were created for the experimental stimuli and an additional 10 for practice. For each room, a second version (its “similar room match”) was created with a 30-degree viewpoint rotation (half rotated clockwise and half rotated counterclockwise). This “similar room match” had the same spatial layout of furniture and wall angles, but with altered visual content: wall colors were changed, and furniture was replaced with different exemplars of the same category (e.g., a table was replaced with a different table). An additional 10 rooms and their altered versions were created for a practice run of the task.

Paintings were chosen from the Google Art Project (https://artsandculture.google.com/). 80 artists were selected, and two paintings were chosen from each artist. The two paintings by each artist (the first painting and its “similar art match”) were similar in terms of style (e.g., choice and use of color, level of detail, brushstrokes) but not necessarily content. An additional set of paintings (10 artists/20 paintings) were selected for practice. None of the practice stimuli overlapped with the experimental stimuli. All stimuli were presented using Psychophysics Toolbox 3 in Matlab (http://psychtoolbox.org/).

### Design and Procedure

The design and procedure were modified from Aly & Turk-Browne (2016a, 2016b). The main difference is that only two images were presented on each trial, rather than five. This was to reduce working memory demands, given the known impairments of hippocampal lesion patients on working memory tasks that require relational representations (e.g., Hannula, Tranel, & Cohen, 2006; Olson et al., 2006; see Yonelinas, 2013 for review).

A stimulus set of 480 unique images was generated such that each of the 160 rooms (80 original rooms and 80 similar room matches) were paired with 3 paintings, all from different artists (likewise, each of the 160 paintings [80 original paintings and 80 similar art matches] were paired with 3 different rooms).

From this stimulus set, 80 image groupings of 6 images each were created (one image grouping for each of the 80 trials in the main experiment). For each trial, one image was selected as the “base image” (the first image presented on that trial). The remaining images were ones that could potentially be presented as the second (comparison) image on that trial **(Figure 2**); a “similar art match” (a room that contained a different painting that was painted by the same artist as that in the base image); a “similar room match” (a room with the same spatial layout as the base image, from a different perspective); an “identical art match” (a room that contained a painting that was identical to that in the base image), an “identical room match” (a room that was identical to the base image); and a non-matching image (an image with a painting by a different artist and a room with a different layout). An image that was an art match (whether identical or similar) to the base image could not also be a room match (whether identical or similar), and vice versa. The same logic followed for an additional 10 image groupings created for a practice run of the task (10 trials).

The 80 trials were split into two tasks: 40 “art” trials and 40 “room” trials. Half of those trials were “identical” trials and half were “similar” trials. Two images were presented on each trial. “Identical” trials involved the presentation of a base image and one of the following: an image with an identical painting (“identical art match”), an image with an identical room (“identical room match”), or a non-matching image. “Similar” trials involved the presentation of a base image and one of the following: its similar art match (different painting by the same artist), its similar room match (room with the same layout from a different perspective), or a non-matching image. Performance on these “similar” trials was hypothesized to benefit from the relational representations of the hippocampus (Eichenbaum & Cohen, 2014).

We define relational representations as representations of multidimensional components of an experience and their unique relationship to one another, such as the relationship between visual features or spatial layouts. The individual features of a given relational representation may be present in other representations as well, but the unique way they combine defines the particular experience. In the current study, performance on similar art trials might benefit from relational representations because individual features of a painting (e.g., use of color, type of brushstroke, level of detail) are not diagnostic of a specific artist. Instead, performance may be helped by representing how those visual features holistically combine to determine an artist’s unique style. Performance on similar room trials would benefit from relational representations because individual features of a room (e.g., the types of furniture, presence of a particular wall angle) are not sufficient to identify configurally similar rooms. Instead, performance on this task requires assessments of furniture layouts and their relationship to the spatial arrangement of wall angles. Thus, accurate performance requires representing the unique relations among pieces of furniture and walls. We note, however, that similar art trials may require less relational processing than similar room trials and, indeed, may potentially be performed reasonably accurately without relational representations. We will return to this issue in the Discussion.

Performance on “identical” trials required even less, or no, relational processing: Instead, accurate performance on these trials can be accomplished by detecting repetitions of particular objects or colors.

On each trial, participants first viewed a cue, either “ART” or “ROOM” for 0.5 s **(Figure 2A)**. This cue instructed the participant to attend to either the style of the paintings (ART) or the layout of the rooms (ROOM). Following this cue, participants viewed a base image for 2.0 s, followed by a 0.5 s interstimulus interval, and then a second image for 2.0 s. The second image could be either an identical art match, an identical room match, a similar art match, a similar room match, or a non-matching image **(Figure 2B)**. Finally, a probe was presented, either “ART?” or “ROOM?”. The probe stayed on the screen until the participant responded; there were no specific instructions to respond as fast as possible. Participants were instructed to respond “yes” if they thought there was a match in the probed dimension and “no” if they thought there was not a match. Responses were made with the 1 and 2 keys, respectively. Participants were to respond “yes” to an “ART?” probe if the two paintings were identical *or* if they were painted by the same artist (identical art match or similar art match), and “no” otherwise. Participants were to respond “yes” to a “ROOM?” probe if the two rooms were identical *or* if they had the same spatial layout from a different perspective (identical room match or similar room match), and “no” otherwise.

For each task (art or room) and trial type (identical or similar), the probability that the attentional cue at the beginning of the trial matched the probe at the end was 80% (valid trials). On the remaining 20% of trials, the cue at the beginning of the trial did not match the probe at the end (invalid trials); Participants were told to attend to one feature (e.g., “ART”) and were probed about whether there was a match on the other feature (e.g., “ROOM?”). The purpose of invalid trials was to ensure that attention was engaged by the cue at the beginning of the trial: if so, participants should be better on valid vs. invalid trials (Posner, 1980).

On any given trial, the second (comparison) image could fall into one of three categories: (1) a cued match (i.e., an art match [either identical or similar] on a trial with an art cue; a room match [either identical or similar] on a trial with a room cue); (2) a non-cued match (i.e., an art match [either identical or similar] on a trial with a room cue; a room match [either identical or similar] on a trial with an art cue); or (3) a non-matching image (neither an art match nor a room match).

On valid trials, the cued (and probed) match was shown 50% of the time, the non-cued (and not probed) match was shown 25% of the time, and a non-matching image was shown the remaining 25% of the time (hence, the correct answer was “yes” half the time and “no” half the time).

On invalid trials, the probed (but not cued) match was shown 50% of the time, the cued (but not probed) match was shown 25% of the time, and a non-matching image was shown the remaining 25% of the time (hence, the correct answer was “yes” half the time and “no” half the time).

The task was blocked: Participants completed 10 trials of a given attentional state before switching to the other (e.g., 10 trials with “ART” cues; 10 trials with “ROOM” cues, and so on). Half of the participants started with art attention and half started with room attention. Similar trials and identical trials were intermixed so that participants could not adopt different strategies during the viewing of the first image of each trial. Specifically, intermixing trials in this way prevents a strategy on identical trials in which participants can simply focus on one feature (e.g., the color of the wall, a corner of a painting) and look for an identical repetition of that feature. Thus, participants had to attend to the entire painting or room on each trial and attempt to extract multidimensional information because it was unknown whether the trial type was similar or identical.

Participants first received instructions and were shown examples of all the different match types (identical art match, identical room match, similar art match, similar room match). Next, they completed 10 practice trials. Participants were required to perform at 80% accuracy to continue to the full experiment. Each person who was tested met this criterion and completed the full task. During the practice and full experiment, participants received feedback after a block of trials (“Wow! You are doing amazingly well! Keep it up!”, “You are doing very well! Keep it up!”, “You are doing ok! Keep it up!”, “This task is challenging, but keep trying!”), as well as the percentage of correct responses. For the practice, they received feedback after every 5 trials; for the full experiment, they received feedback every 10 trials.

### Statistical Analyses

Analysis of variance (ANOVAs) and follow-up t-tests were conducted in Matlab. All reported p-values are two-tailed, with values less than 0.05 considered statistically significant. 95% confidence intervals are reported where appropriate. Effect sizes (partial eta squared [*η* _p_^2^] for ANOVAs and Cohen’s *d*_s_ and *d*_z_ for t-tests) were implemented following Lakens (2013). Stimuli, code, and data can be found on GitHub: https://github.com/alylab/artmusePatient.

## Results

We analyzed behavioral sensitivity (A’; 1 = perfect, 0.5 = chance; Donaldson, 1992) and response times (RTs) to the “ART?” and “ROOM?” probes. A’ was chosen as the measure of behavioral sensitivity because it is non-parametric and because it is the measure we have used in prior studies with a similar task (Aly & Turk-Browne 2016a; Aly & Turk-Browne 2016b). Log-transformed RTs were used in all analyses, given that raw RT values can be highly skewed.

### Valid vs. Invalid trials

We first examined whether attention was effectively guided by the cue at the beginning of the trial. If so, participants should be more accurate and faster on valid vs. invalid trials. To that end, we conducted four separate 3-way repeated-measures ANOVAs, two for patients and two for healthy participants. The dependent measures were A’ and RTs. The independent variables were condition (similar vs. identical trials), attentional state (art vs. room trials), and validity (valid vs. invalid trials).

For A’, there was a significant main effect of condition for healthy participants [*F*_1,13_ = 8.68, *p* = 0.01, η_p_^2^ = 0.40] reflecting better performance on identical vs. similar trials (this effect was marginally significant for patients [*F*_1,6_ = 5.83, *p* = 0.052, *η* _p_^2^ = 0.49]). There was also a significant condition by attentional state interaction for healthy participants [*F*_1,13_ = 7.18, *p* = 0.019, η_p_^2^ = 0.36]; Collapsing across trial validity, performance was better on identical room vs. identical art trials, but better on similar art vs. similar room trials. Importantly, there was a significant main effect of validity for both the healthy participants [*F*_1,13_ = 34.00, *p* = 0.00006, *η* _p_^2^ = 0.72] and the patients [*F*_1,6_ = 33.82, *p* = 0.0011, *η* _p_^2^ = 0.85], indicating that behavioral sensitivity was better on valid vs. invalid trials. No other main effects or interactions were significant [patients: all other *p*s > 0.16; healthy participants: all other *p*s > 0.094]. Moreover, A’ on invalid trials was not different from chance for any trial type, for either healthy participants or patients (all *p*s > 0.09).

For RT, there was a significant main effect of validity for the healthy participants [*M_valid_*= 1.53s, *M_invalid_* = 2.34s; *F*_1,13_ = 27.39, *p* = 0.0002, *η* _p_^2^ = 0.68], indicating faster RTs on valid vs. invalid trials. This comparison did not reach statistical significance for the patients [*M_valid_* = 1.21s, *M_invalid_* = 2.25s; *F*_1,6_ = 4.46, *p* = 0.08, *η* _p_^2^ = 0.43]. No other main effects or interactions were significant [patients: all other *p*s > 0.15; healthy participants: all other *p*s > 0.10].

Taken together, these results suggest that attention was effectively engaged by the cue at the beginning of the trial: Healthy participants were more accurate and faster on valid vs. invalid trials. Patients were more accurate on valid vs. invalid trials. Furthermore, performance was not different from chance on invalid trials for either patients or healthy participants, indicating a particularly strong manipulation of attention. Having confirmed that attention was effectively modulated, we next focus analyses on valid trials.

### Identical Trials

We examined A’ and RTs on identical trials with 2 (attentional state; art or room) by 2 (group: patient or healthy participant) mixed-model ANOVAs **(Figure 3A)**. For A’, there was no main effect of attentional state [*F*_1,19_ = 3.66, *p* = 0.07, *η* _p_^2^ = 0.16], no main effect of group [*F*_1,19_ = 3.19, *p* = 0.09, *η* _p_^2^ = 0.14], and no group by attentional state interaction [*F*_1,19_ = 0.08, *p* = 0.78, *η* _p_^2^ = 0.004]. Similarly, for RTs, there was no main effect of attentional state [*F*_1,19_ = 0.10, *p* = 0.75, *η* _p_^2^ = 0.01], no main effect of group [*F*_1,19_ = 3.03, *p* = 0.10, *η* _p_^2^ = 0.14], and no group by attentional state interaction [*F*_1,19_ = 1.10, *p* = 0.31, *η* _p_^2^ = 0.05]. Thus, performance was well-matched on identical art and identical room trials, and there was no statistically significant difference between patients and healthy participants.

**Figure 3.**
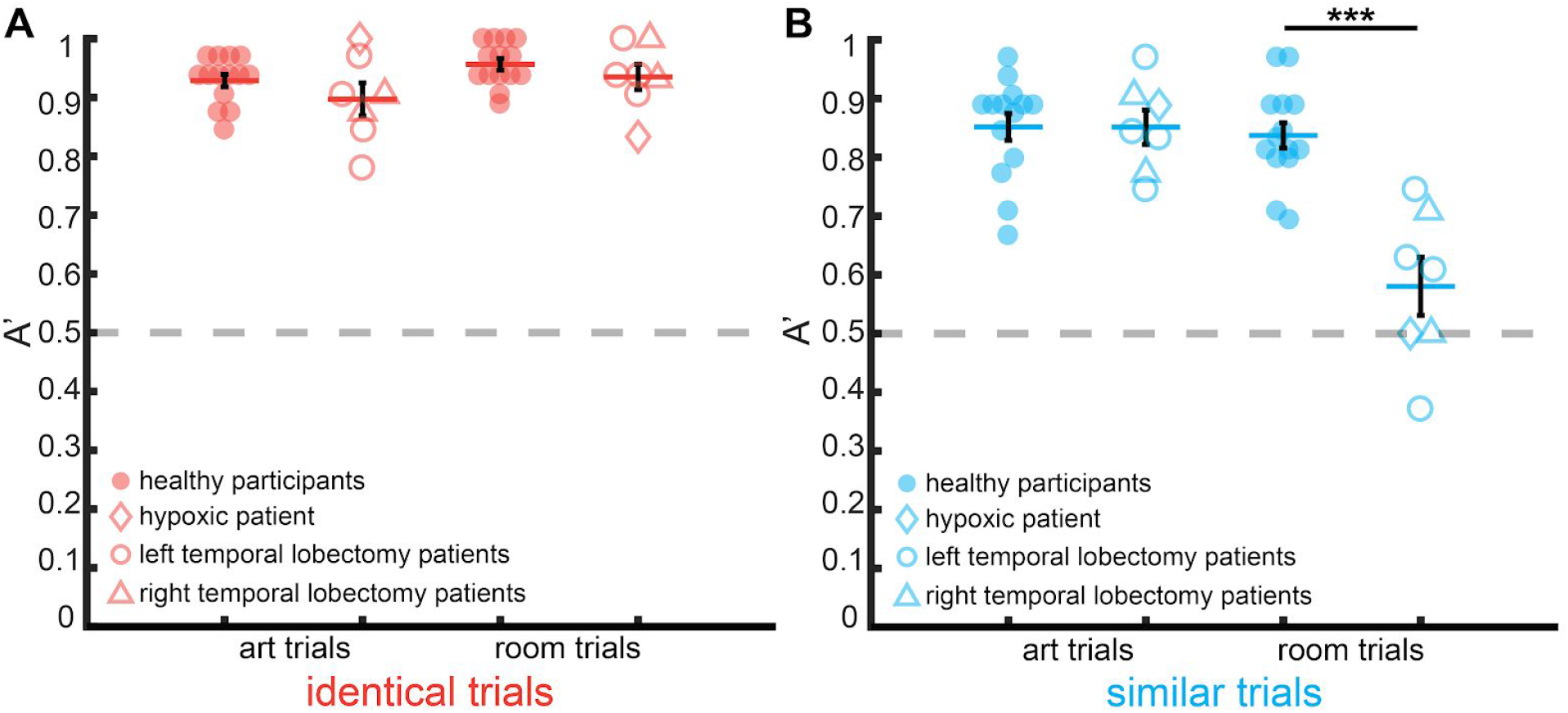
Behavioral performance (A’). **(A)** There was no statistically significant difference between healthy participants and patients on identical art trials or identical room trials. **(B)** There was no statistically significant difference between patients and healthy participants on similar art trials, but the patients were significantly impaired on similar room trials — where their average performance was no higher than chance (dashed line). Error bars indicate standard error of the mean. *** *p* <.0001.

Performance on identical trials was quite high overall. Thus, one concern is that the patients may not be impaired because the task was simply too easy — i.e., that there were ceiling effects. However, an individual patient did perform significantly worse than age-matched healthy individuals on identical room trials (see *Analyses by Patient Group*), indicating that this task was sensitive to behavioral impairments. Furthermore, the analysis of similar trials (discussed below) demonstrates that task difficulty cannot explain the pattern of results observed for the patients.

### Similar Trials

We next examined A’ and RTs on similar trials using the same 2 (attentional state: art or room) by 2 (group: patient or healthy participant) mixed-model ANOVAs **(Figure 3B)**. For RTs, there was no main effect of attentional state [*F*_1,19_ = 0.80, *p* = 0.38, *η* _p_^2^ = 0.04], no main effect of group [*F*_1,19_ = 3.63, *p* = 0.07, *η* _p_^2^ = 0.16], and no group by attentional state interaction [*F*_1,19_ = 2.53, *p* = 0.13, *η* _p_^2^ = 0.12].

For A’, there was a main effect of attentional state [*F*_1,19_ = 21.85, *p* = 0.0002, *η* _p_^2^ = 0.53], a main effect of group [*F*_1,19_ = 19.16, *p* = 0.0003, *η* _p_^2^ = 0.50], and a significant group by attentional state interaction [*F*_1,19_ = 17.80, *p* = 0.0005, η_p_^2^ = 0.48]. Given this interaction, we conducted follow-up *t*-tests to compare A’ values for healthy participants and patients on similar art and similar room trials.

Patients and healthy participants were not different on similar art trials [*t*_19_ = −0.004, *p* = 1.0, 95% CIs = −0.08 – 0.08, Cohen’s *d*_s_ = 0.002]. However, patients were significantly impaired, relative to healthy participants, on similar room trials [*t*_19_ = −5.66, *p* = 0.00002, 95% CIs = −0.35 – −0.16, Cohen’s *d*_s_ = 2.62]. This selective impairment was not a result of differing task difficulty for similar room vs. similar art trials because healthy participants performed just as well on both trial types [*t*_13_ = −0.43, *p* = 0.67, 95% CIs = −0.08 – 0.06, Cohen’s *d*_z_ = 0.12]. Patients, however, performed significantly worse on similar room vs. similar art trials [*t*_6_ = −4.70, *p* = 0.0033, 95% CIs = −0.41 —0.13, Cohen’s *d*_z_ = 1.78], and in fact their performance on similar room trials was not different from chance [*t*_6_ = 1.64, *p* = 0.15, 95% CIs = 0.46 – 0.70; chance = 0.5]. Patients were significantly above chance (all *p*s < 0.0001) on all other trial types (identical art, identical room, similar art: healthy participants performed above chance on every trial type, all *p*s < 0.0001).

To confirm that this pattern of results is not contingent on the chosen measure of behavioral performance (i.e., A’), we re-did the analyses with accuracy and d’ as the behavioral measures of interest and replicated the same pattern of results. Note that for d’, the Snodgrass & Corwin (1988) correction was applied to avoid hit rates of 1 and false alarm rates of 0.

Specifically, for accuracy, patients were not different from healthy participants on identical art trials (*t*_19_ = −1.28, *p* = 0.22, 95% CIs = −0.16 – 0.04, Cohen’s *d*_s_ = 0.59), identical room trials (*t*_19_ = −0.89, *p* = 0.39, 95% CIs = −0.10 – 0.04, Cohen’s *d*_s_ = 0.41), or similar art trials (*t*_19_ = 0.10, *p* = 0.93, 95% CIs = −0.09 − 0.10, Cohen’s *d*_s_ = 0.04). However, they were significantly impaired on similar room trials (*t*_19_ = −3.98, *p* = 0.0008, 95% CIs = −0.29 - −0.09, Cohen’s *d*_s_ = 1.84).

For d’, patients were not different from healthy participants on identical art trials (*t*_19_ = −0.99, *p* = 0.34, 95% CIs = −0.82 - 0.29, Cohen’s *d*_s_ = 0.46), identical room trials (*t*_19_ = −0.85, *p* = 0.40, 95% CIs = −0.82 − 0.34, Cohen’s *d*_s_ = 0.40), or similar art trials (*t*_19_ = −0.04, *p* = 0.97, 95% CIs = −0.60 - 0.58, Cohen’s *d*_s_ = 0.02). However, they were significantly impaired on similar room trials (*t*_19_ = −4.32, *p* = 0.0004, 95% CIs = −1.62 - −0.56, Cohen’s *d*_s_ = 2.00).

The measures of A’ and d’ include both hits and false alarms. Thus, it is unclear from the above results whether the patients’ impairment on similar room trials is a result of a reduced hit rate, an increased false alarm rate, or both. We therefore compared the hit and false alarm rates for patients and healthy participants. Patients had significantly reduced hit rates on similar room trials, relative to healthy participants [*t*_19_ = −2.88, p = 0.0095, 95% CIs = −0.54 - −0.09, Cohen’s *d*_s_ = 1.33], but false alarm rates did not differ [*t*_19_ = 0.78, p = 0.44, 95% CIs = −0.10 - 0.23, Cohen’s *d*_s_ = 0.36]. There was no difference between patients and healthy participants in hit or false alarm rates on any other trial type (all *p*s > 0.13).

Taken together, patients were significantly and selectively impaired on similar room trials, and their performance was not different from chance.

### Comparison of Identical and Similar Trials

Patients showed statistically significant impairment only on similar room trials. However, on identical trials, the main effect of group (patients vs. healthy participants) approached significance [*F*_1,19_ = 3.19, *p* = 0.09, *η* _p_^2^ = 0.14]. We therefore directly compared identical and similar trials to test whether patients were significantly more impaired on similar trials, but only when attention was directed to room layouts. To this end, we conducted two 2 (group: patient or healthy participant) by 2 (condition: identical or similar trials) mixed-model ANOVAs, one for art trials and one for room trials.

For art trials, there was a main effect of condition [*F*_1,19_ = 9.21, p = 0.007, *η* _p_^2^ = 0.33], indicating better performance on identical vs. similar trials. However, there was no main effect of group [*F*_1,19_ = 0.39, p = 0.54, *η* _p_^2^ = 0.02] nor a group by condition interaction [*F*_1,19_ = 0.57, p = 0.46, *η* _p_^2^ = 0.03]. Thus, there was no statistically significant difference between the performance of patients and healthy participants on identical art trials or similar art trials.

For room trials, there was a main effect of condition [*F*_1,19_ = 103.36, p < 0.00001, *η* _p_^2^ = 0.84], indicating better performance on identical vs. similar trials. There was also a main effect of group [*F*_1,19_ = 27.97, p = 0.00004, *η* _p_^2^ = 0.59] and a group by condition interaction [*F*_1,19_ = 25.37, p = 0.00007, η_p_^2^ = 0.57]. This result indicates that patients were significantly more impaired on similar room trials than identical room trials. Indeed, comparison of patients and healthy participants on identical room trials revealed no statistically significant difference [*t*_19_ = −1.05, p = 0.31, 95% CIs = −0.06 - 0.02, Cohen’s *d*_s_ = 0.49].

### Analyses by Patient Group

The above analyses include temporal lobectomy patients and the hypoxia patient. However, the hypoxia patient is less well-characterized than the lobectomy patients: Fewer neuropsychological test scores were available, and the extent of brain damage is less clear in the hypoxia patient vs. the lobectomy patients. To ensure that the results are not unduly driven by the hypoxia patient, we analyzed the groups separately.

First, we compared the hypoxia patient to healthy participants (n=4) who were matched in age (patient 101: 19 years old, healthy participants: *M_age_*= 20.00 years) and education (patient 101; 12 years, healthy participants: *M_education_*= 14 years). To conduct this analysis, we used a method derived by Crawford and Howell (1998; also see Crawford, Garthwaite, & Howell, 2009) that allows for the comparison of a single patient against a sample of healthy participants. Patient 101’s performance (A’) did not differ from that of healthy participants on art trials [identical art: *t*_3_ = 1.63, *p* = 0.90, 95% CIs = −0.05 – 0.16; similar art: *t*_3_ = −0.48, *p* = 0.33, 95% CIs = −0.17 – 0.13]. However, he was significantly impaired on both types of room trials [identical room: *t*_3_ = −3.35, *p* = 0.02, 95% CIs = −0.22 – −0.006; similar room: *t*_3_ = −4.37, *p* = 0.01, 95% CIs = −0.70 —0.11]. Nevertheless, his performance on identical room trials was well above chance (0.83), while his performance on similar room trials was exactly at chance (0.50). Thus, the hypoxia patient exhibited general impairments in spatial processing, and his performance on spatial relational trials (i.e., similar room trials) was at chance.

The temporal lobectomy patients exhibited the same pattern of results as the main group analysis. They were significantly impaired on similar room trials relative to age- and education-matched healthy participants, but performed normally on all other trial types [identical art: *t*_14_ = −1.58, *p* = 0.14, 95% CIs = −0.10 – 0.01, Cohen’s *d*_s_ = 0.82; identical room: *t*_14_ = −0.44, *p* = 0.67, 95% CIs = −0.05 – 0.03, Cohen’s *d*_s_ = 0.23; similar art: *t*_14_ = 0.38, *p* = 0.71, 95% CIs = −0.08 – 0.11, Cohen’s *d*_s_ = 0.20; similar room: *t*_14_ = −4.37, *p* = 0.0006, 95% CIs = −0.32 – −0.11, Cohen’s *d*_s_ = 2.26]. As with the hypoxia patient, the temporal lobectomy patients were not above chance on similar room trials [*t*_5_ = 1.68, *p* = 0.15, 95% CIs = 0.45 – 0.74; chance = 0.5].

Thus, the main results are not driven by the inclusion of the hypoxia patient. Indeed, the selectivity of behavioral deficits on similar room trials is more apparent in the temporal lobectomy patients. The hypoxia patient exhibited a more general impairment in spatial discrimination (i.e., impairment on both identical and similar room trials). This might have occurred because only the hypoxia patient had bilateral hippocampal damage (the temporal lobectomy patients had unilateral damage). Alternatively, the hypoxia patient might have shown a more general impairment in spatial processing because of damage outside of the hippocampus/MTL (Guderian et al., 2015; Huang & Castillo, 2008).

It is worth noting that one left hemisphere temporal lobectomy patient had quite widespread brain damage extending to the frontal lobes. The remaining patients had damage confined to the temporal lobe. To ensure that the inclusion of this frontal lesion patient is not unduly influencing the results, we reanalyzed the data with this patient excluded. We first confirmed that patients and healthy participants were still matched in age and education without this patient, and that was indeed the case (age: *t*_18_ = 0.02, *p* = 0.99, 95% CIs = −19.00 − 19.34; education: *t*_18_ = −0.12, *p* = 0.91, 95% CIs = −2.25 − 2.01). The main results held in this analysis. Specifically, the patients were selectively and significantly impaired on similar room trials (identical art: *t*_18_ = −1.24, *p* = 0.23, 95% CIs = −0.09 - 0.02; Cohen’s *d*_s_ = 0.60; identical room: *t*_18_ = −1.58, *p* = 0.13, 95% CIs = −0.08 - 0.01; Cohen’s *d*_s_ = 0.77; similar art: *t*_18_ = 0.47, *p* = 0.65, 95% CIs = −0.07 - 0.10; Cohen’s *d*_s_ = 0.23; similar room: *t*_18_ = −6.29, *p* =.000006, 95% CIs = −0.38 - −0.19; Cohen’s *d*_s_ = 3.07).

Thus, the observed results are not driven by the hypoxia patient or the patient with frontal cortex damage. Because all patients had hippocampal/MTL damage, and damage outside of these regions varied across patients, it is likely that MTL damage either directly or indirectly led to the observed behavioral deficits on the similar room task. We return to this issue in the Discussion.

### Impairment of Attention or Perception?

Our goal in this study was to determine if hippocampal/MTL damage impaired attentional behavior. The impairment on similar room trials, however, is consistent with either an impairment in attention, an impairment in perception, or both. That is, it could be the case that patients fail to attend to task-relevant features that would enable them to identify relationally similar rooms. Alternatively, they may attend to these features but cannot perceive them properly. A key marker of successful attention is performance enhancements on valid vs. invalid trials (Posner, 1980), which offers a potential way of disentangling perception and attention deficits. We therefore conducted a follow-up analysis on similar room trials to determine whether patients exhibit a reduced benefit of valid cueing relative to healthy participants. If so, this would be consistent with a deficit in attention. If, however, patients benefit just as much from valid cueing as healthy participants, their impairment would be more in line with a deficit in perception. We note in advance that this analysis is expected to be noisy because there are only 4 invalid trials per participant in the similar room task. But we conducted it nevertheless, as a follow-up analysis that may be useful in inspiring future studies.

We carried out a 2 (group: healthy participant or patient) by 2 (validity: valid or invalid) mixed model ANOVA on A’ for similar room trials, looking specifically for an interaction. The group by validity interaction did not reach significance (*F*_1,19_ = 2.40, p = 0.14, *η* _p_^2^ = 0.11). Nevertheless, as an exploratory analysis, we examined the valid vs. invalid benefit for healthy participants and patients with separate t-tests. Healthy participants showed robust improvements in A’ on valid vs. invalid similar room trials (*t*_13_ = 5.64, *p* =.00008, 95% CIs = 0.28 - 0.64; Cohen’s *d*_z_ = 1.51). In contrast, patients did not show statistically significant improvement in A’ on valid vs. invalid similar room trials (*t*_6_ = 2.03, *p* =.09, 95% CIs = −0.05 - 0.53; Cohen’s *d*_z_ = 0.77). It is worthwhile to note that, although the interaction failed to reach significance, the validity effect size for healthy participants was nearly twice that for patients (*d_z_* = 1.51 vs *d_z_ =* 0.77, respectively).

While these exploratory analyses hint at an attentional impairment, they are not strong enough for us to claim that we have identified an attention impairment rather than a perceptual one. Future studies with larger sample sizes, and more invalid trials, will be necessary to definitively test for an attentional deficit in MTL amnesics. We return to this issue in the Discussion.

## Discussion

### Summary

We examined whether the hippocampus/MTL critically support attention and perception, and what their contribution might be. We tested individuals with hippocampal/MTL damage and healthy age- and education-matched participants on attention tasks that varied in their demands on relational and spatial processing. We found that both spatial and relational representations were important components of MTL contributions: Patients with hippocampal/MTL damage were selectively impaired on spatial discriminations, and spatial *relational* discriminations in particular.

These results provide evidence that the hippocampus plays a critical role in rapid perceptual processes that benefit from attention. They add to a growing body of literature highlighting the far reach of the hippocampus in cognition, including perception (Lee et al., 2012), working memory (Yonelinas, 2013), implicit memory (Hannula & Greene, 2012), decision making (Shohamy & Daw, 2015), imagination (Schacter et al., 2017), creativity (Rubin et al., 2015), language (Duff & Brown-Schmidt, 2012), and social cognition (Schafer & Schiller, 2018). These findings, and our new results, together pose a strong challenge to theories of hippocampal function that view it as a system that is dedicated for long-term memory (Squire & Wixted, 2011).

### Relation to Prior Work

Although we have emphasized the attentional demands of our task, the task also placed demands on perception. Indeed, we believe it is very difficult (or impossible) to study attention separately from perception, because the key behavioral marker of attention is improvements in perceptual behavior. We nevertheless conducted some exploratory analyses to determine whether patients’ impairments on similar room trials could be attributed to a failure to deploy attention properly vs. a failure to perceive the relevant features. If attention is deployed effectively, performance should be substantially better on trials with valid attention cues vs. those with invalid cues (Posner, 1980). We found that patients did not show statistically significant improvements in performance on valid vs. invalid similar room trials. Moreover, on similar room trials, the effect size for performance enhancements on valid vs invalid trials was nearly doubled for healthy participants relative to patients. Nevertheless, because we did not observe a statistically significant interaction between group (patient or healthy participant) and cue validity (valid vs invalid) on similar room trials, we are cautious in overly interpreting these results. Future studies will be needed to explore this issue more fully, with analyses that are more highly powered. Thus, we conclude that the impairment that patients have demonstrated on similar room trials might be one of attention, perception, or interactions between the two.

Our findings therefore complement — and extend — studies on perception in hippocampal amnesia (Lee et al., 2012; Yonelinas, 2013) in several ways. For example, studies of perception often find that patients with relatively selective hippocampal damage are impaired on tasks that use scenes as stimuli, and not those that use faces, objects, art, or colored shapes (Barense et al., 2005, 2007; Behrmann et al., 2016; Graham et al., 2006; Lee et al., 2005a, 2005b; but see Erez et al., 2013; Warren et al., 2010, 2011; also see Goodrich & Yonelinas, 2016). However, perceptual impairments for non-scene stimuli can be observed in patients with hippocampal damage, primarily when the task requires relational processing (e.g., Warren et al., 2011, 2012). That both spatial and relational processing are important (Aly et al., 2013; Aly & Turk-Browne, 2018; Hannula, Tranel, & Cohen, 2006) is supported by findings that scene perception impairments in hippocampal lesion patients are pronounced when the task involves changes in perspective, increasing the demands on relational processing (e.g., Behrmann et al., 2016; Erez et al., 2013; Lee et al., 2005b; also see King et al., 2002).

Here, we find converging evidence that spatial and relational processing are both critical features of hippocampal/MTL function: Discrimination of spatial features was more impaired than discrimination of artistic features, but only when relational processing demands were high. This result is noteworthy because the temporal lobectomy patients also had extensive damage to the perirhinal cortex, which could have impaired attention to, and perception of, artistic features in the similar art task (c.f., Lee et al., 2005a). One potential reason why patients did not show impairment on the similar art task is that it might have been possible to solve it on the basis of basic or broad categorical features (e.g., spatial frequency content or color across paintings) without forming a relational or conjunctive representation. We return to this issue below, in *Future Directions*.

Our results go beyond other studies of perception in a number of ways. First, we presented stimuli for a relatively brief amount of time and with a very short inter-stimulus interval, whereas many studies of perception in patients with MTL lesions present stimuli for a longer duration and/or with a longer interval between stimulus presentations. For example, several studies present images until the participant responds (e.g., Barense et al., 2007; Behrmann et al., 2016; Erez et al., 2013; Graham et al., 2006; Lee et al., 2005b; Hartley et al., 2007; Warren et al. 2011) and/or require individuals to remember a target stimulus across many trials (e.g., Barense et al., 2005; Lee et al., 2005a). Although such paradigms convincingly tax perception and the results from them are difficult to explain solely in terms of long-term memory impairments, these paradigms nevertheless leave room for attention, perception, working memory, and/or long-term memory to interact to contribute to performance. Here, we attempted to minimize the influence of working- and long-memory and tax rapidly evolving attention and perception (e.g., Mullally, Intraub, & Maguire, 2012) by using relatively brief stimulus presentations, very short inter-stimulus intervals, and trial-unique images.

Second, we manipulated participants’ attentional states (e.g., attention to artistic style vs. room layouts) while holding the type of stimulus constant (also see Hartley et al., 2007). In contrast, most studies of perception in patients with MTL damage investigate different kinds of perception by varying the stimulus itself (e.g., presenting paintings vs. scenes on different trials: Lee et al., 2005a).

Finally, we designed the similar image tasks, particularly the similar room task, so that low-level visual features were not particularly useful for task performance: abstract relations were needed to accurately guide behavior. In contrast, in some studies of perception in MTL amnesics, feature- or item-level information can be sufficient for task performance, even if the intention is to only manipulate relational information (see Aly et al., 2013; Baxter, 2009). The approach that we took for similar image trials (particularly similar room trials) — preserving relational information across images that varied in low-level visual features (also see Hartley et al., 2007) — is complementary to approaches taken in some studies of perception, where relational information is manipulated across images that are otherwise identical in low-level visual features (e.g., Aly et al., 2013; Behrmann et al., 2016; Erez et al., 2013; Lee et al., 2005b).

### Limitations of Patient Studies

We have emphasized the hippocampal and medial temporal lobe lesions in the patients involved in the current study. However, it is important to note that temporal lobe epilepsy is associated with abnormalities in, and reduction in gray matter throughout, widespread brain networks (e.g. Bonilha et al., 2004). While temporal lobectomy surgical lesions can be relatively focal, the brains of these patients may have disrupted functioning in regions anatomically or functionally connected to the hippocampus and medial temporal lobe cortex.

Likewise, hippocampal damage (from etiologies beyond temporal lobe epilepsy) can cause broader network abnormalities. Even in cases of rather focal hippocampal atrophy, volumetric changes to the extended hippocampal network, such as the entorhinal cortex and thalamus, can be observed (Argyropoulos et al., 2019). Functional abnormalities have also been noted, e.g., decreases or changes in functional connectivity, including between non-hippocampal regions, and changes in overall activity outside of the hippocampus (Argyropoulos et al., 2019; Henson et al., 2016).

As with any lesion study, it is therefore difficult to determine exactly which aspect of disrupted brain functioning is key for the observed behavioral impairments. However, because all of the patients had hippocampal/MTL damage — and damage outside of the MTL varied across patients — it is likely that this damage either directly or indirectly (through its effects on other brain regions) is an important determinant of the behavioral deficits. That said, the mechanisms by which these behavioral impairments arise is not clear from the current study. There are at least two possibilities: (1) the hippocampus/MTL is involved in assessing relational similarities between scenes, or (2) the hippocampus/MTL provides important input or output to other regions that are involved in assessing relational similarities between scenes. Functional neuroimaging studies of patients with hippocampal lesions will be informative in this regard: Such studies can illuminate how broader network function might be disrupted in these patients while they are performing the attention tasks used here. This would shed light on how and why hippocampal/MTL damage impairs performance. Nevertheless, we can conclude that the hippocampus/MTL is critical for spatial relational attention and/or perception, even if the mechanisms by which this happens are not yet clear.

Finally, it is worth noting that the temporal lobectomy patients had unilateral hippocampal lesions. As a result, some memory functions — as assessed by neuropsychological testing — are only mildly rather than severely impaired. Thus, the perceptual deficits these patients exhibit in the current study are notable: One could have predicted that they would not have been impaired due to the remaining, intact hippocampus/MTL. The current results therefore suggest that unilateral MTL damage may be sufficient for impairments in online spatial relational discriminations. Relational attention tasks such as the one used here may be particularly sensitive to reductions in MTL integrity.

### Future Directions

The evidence reported here is a valuable contribution to the literature given the difficulty of finding, characterizing, and testing patients with medial temporal lobe damage. Many studies of amnesic patients test only a few individuals, given the rarity of the population of interest. Our sample size is comparable to, or larger than, studies investigating perceptual impairments in patients with medial temporal lobe damage (e.g., Aly et al., 2013; Barense et al., 2005, 2007; Behrmann et al., 2016; Erez et al., 2013; Graham et al., 2006; Hartley et al., 2007; Lee et al., 2005a, 2005b; Warren et al., 2010, 2011, 2012). As for any patient study, however, generalizing our results by testing more patients with medial temporal lobe damage — particularly selective hippocampal lesions — will be important. This is especially the case because the hypoxia patient, unlike the patients with temporal lobectomies, also exhibited an impairment on spatial attention trials that did not place heavy demands on relational processing. Thus, one possibility is that bilateral hippocampal damage can produce impairments in attention and/or perception when spatial representations are required, even if relational processing demands are minimal. An alternative possibility is that damage outside of the MTL in the hypoxia patient contributed to a more general impairment. Future studies involving patients with confirmed selective hippocampal lesions will be important.

The patient results reported here converge with our previous fMRI studies, which demonstrated that the hippocampus is more strongly modulated by attention to spatial layouts vs. artistic features, and predicted behavior most strongly for spatial relational attention (Aly & Turk-Browne, 2016a; Aly & Turk-Browne, 2016b). Likewise, we found that MTL cortical regions were more strongly modulated by attention to spatial relations (Aly & Turk-Browne 2016a; Aly & Turk-Browne, 2016b). The patient and neuroimaging findings therefore converge in showing an important role for the MTL in attention tasks, particularly those that assess attention to, and perception of, spatial relational features.

One limitation of the current study, and our prior fMRI work (Aly & Turk-Browne 2016a; Aly & Turk-Browne, 2016b), is that it is possible that the similar art task was simply not as “relational” as the similar room task. In the similar room task, many types of features had to be bound and compared (e.g., angles and lengths of walls, placement and type of furniture). Conversely, in the similar art task, individuals may have chosen to focus on the choice of colors, spatial frequency content, or broad categorical features (e.g., nature scene vs. city scene). None of these strategies would be perfect, because spatial frequency content, choice of color, and scene category were not individually diagnostic of paintings by the same artist. For example, many paintings were of an Impressionist or Post-Impressionist style, so common artistic features were present across paintings by different artists. To perform well, participants would ideally not just attend to the presence or absence of particular color palettes or spatial frequencies, but to the combination of those features. All that said, it is not clear what performance level can be accomplished on the similar art task by attending to individual independent features vs the unique holistic conjunction of those features. This is a weakness relative to other tasks in which the usefulness of feature vs conjunction information was carefully manipulated (e.g., Barense et al., 2005; Bussey et al., 2002). Thus, it is possible that there was less relational processing on similar art vs. similar room trials, and good performance on similar art trials may have been possible without relational representations. For example, individuals may have chosen to treat the paintings as a “unitized” whole rather than in terms of associations between individual features (Mayes, Montaldi, & Migo, 2007). Thus, a key difference between the similar art and similar room tasks may be in the amount of relational processing required, in addition to the requirement to attend to object-based vs. spatial features (or, alternatively, smaller vs. larger parts of the image). However, these tasks were equally difficult for healthy individuals, alleviating the concern that the similar art task was simply less challenging or less complex. The fact that the similar art and similar room tasks were equally difficult for healthy individuals is also key for eliminating the concern that the patients are simply more impaired on harder tasks.

Nonetheless, to comprehensively demonstrate that the hippocampus/MTL is selectively involved in relational attention when such attention taxes spatial representations, other forms of relational attention must be examined (for a similar approach in memory, see Konkel et al., 2008). One promising approach is to investigate whether the hippocampus plays a critical role in attention to temporal relations. In a recent fMRI study (Cordova et al., 2019), we found that the hippocampus is more strongly modulated by attention to temporal vs. size or spatial relations in a rapid, relatively simple stimulus display. A role for the hippocampus in attending to temporal relations would be consistent with a vast literature implicating the hippocampus in temporal and sequential processing (Aly et al., 2018; Barnett et al., 2014; Davachi & DuBrow, 2015; DuBrow & Davachi, 2014; Eichenbaum, 2013; Palombo et al., 2016; Ranganath, 2019; Thavabalasingam et al., 2019). For example, neuropsychological studies indicate that the hippocampus plays an essential role in estimating the temporal duration of events (Palombo et al., 2016; Palombo & Verfaellie, 2017). Thus, one compelling avenue for future research is to determine whether the hippocampus makes a critical contribution to temporal attention (e.g., Cordova et al., 2019; Nobre & van Ede, 2018), which would complement existing studies examining its role in temporal memory.

### Conclusion

The medial temporal lobe makes a critical contribution to rapid perceptual processes that benefit from attention. Damage to this region selectively impairs discrimination of spatial relations. Such an impairment was observed in a task that placed no demands on long-term memory, demonstrating the importance of medial temporal lobe function even on the timescale of online visual processing. This evidence joins a growing body of work highlighting the ubiquity of medial temporal lobe contributions to cognition, contributions that may be realized via flexible, spatial, and relational representations.

## Author Contributions

N.R. performed research, analyzed data, wrote the paper. M.M. performed research, contributed analytic tools. S.A. performed research. M.A. designed research, performed research, analyzed data, wrote the paper.

## Acknowledgments

We would like to thank all the researchers and clinicians of the New York University Patient Registry for the Study of Perception, Emotion, and Cognition (NYU PROSPEC) for their involvement with patient recruitment and patient data collection. We would also like to thank Frank Provenzano, Jeanelle France, and Evie Sobczak for assistance with structural MRI analyses for patient 101, and Eren Günseli for obtaining structural MRI scans for healthy individuals. This work was funded by the following grants (to M.A.); an NSF CAREER Award (BCS-1844241), a Brain & Behavior Research Foundation NARSAD Young Investigator Grant (27893), and a Zuckerman Institute Seed Grant for MR Studies (CU-ZI-MR-S-0001).

